# Synthesizing human-specific spike-in standards for RNA-seq experiments and assessing their technical performance

**DOI:** 10.1101/2025.04.01.646725

**Authors:** Rui Qin, Wei Fan, Feng Ding, Sen Wang, Bing He, Mingming Hou, Qiang Lin, Peng Cui, Wanfei Liu

## Abstract

Using spike-in standards for RNA-seq experiments is critical to evaluate technical bias introduced during sample preparation, sequencing and analysis. Although some external RNA spike-in standards have been developed, species-specific spike-in standard was not reported yet. Here we developed the human-specific spike-in standards with 65 controls. We first extracted human endogenous RNAs with various lengths and GC contents, introduced random mutations approximately every 75 bp in each RNA, and then synthesized these RNAs as spike-in RNAs. After that, four mixtures of these spike-in RNAs covering a 2^20^ dynamic range were obtained. To ensure the accuracy of RNA concentration, two rounds of ddPCR were conducted for each spike-in RNA and the intraclass correlation coefficient between two ddPCRs ranged from 0.9954 to 0.9971 after removing the two spike-in RNAs with the largest concentration difference. Furthermore, we showed that the sequencing error profiles were distinct between platforms and the library preparation procedures were related with the discrepancies in spike-in RNA read percent, transcript abundance, sequence coverage distribution, and differential gene expression. In addition, two regression models of sequence coverage were built based on RNA second structure and GC content, and 86.62%–91.78% of the variation can be explained. Our study demonstrates the technical performance of the human-specific spike-in standards for RNA-seq experiments and illustrates the biases across different libraries, platforms, and laboratories.

## Introduction

RNA-seq provides a far more precise measurement of transcript abundance and has already revolutionized transcriptome studies (Wang et al. 2009; Stark et al. 2019). However, various experimental and sequence-related factors can interfere with the RNA-seq results (Shi et al. 2021). An appropriate normalization methods can reduce the effect of various interfering factors, such as quality control of experimental procedures and standardized methods for data analysis (Conesa et al. 2016). Nevertheless, the bias introduced in RNA-seq is difficult to be accurately evaluated. For instance, a benchmarking study on RNA-seq showed great inter-laboratory variation, mainly due to RNA enrichment, strandedness, and data analysis methods (Wang et al. 2024). Furthermore, Stoler et al. analyzed 1,943 datasets across seven Illumina sequencing platforms, demonstrating significant inter-platform divergence with notable intra-platform fluctuation (Stoler and Nekrutenko 2021). Reference standards have been explored to evaluate the technical performance of next-generation sequencing technologies and to mitigate the negative effects of aforementioned interfering factors (Hardwick et al. 2017).

Evaluation of RNA-seq using internal controls is a feasible method. Endogenous internal controls have been widely used, but the expression of commonly used endogenous reference genes tends to fluctuate in different conditions (Dundas and Ling 2012; Zeng et al. 2016). External RNA controls, employed as a spike-in standard, can be used for quality control and quantification comparisons (Yang 2006). In 2003, the External RNA Controls Consortium (ERCC) was formed, and external RNA controls were developed to evaluate technical performance in microarray and RNA-seq experiments (Baker et al. 2005; Pine et al. 2016). External RNA controls were systematically evaluated based on multiple performance characteristics, such as the limit of detection, signal response to concentration, and differential expression (Jiang et al. 2011; Munro et al. 2014).

Although ERCC has been widely adopted, it may still fail to fully capture the complexity or behavior of natural RNA samples because ERCC is mainly composed of non-human or artificial sequences (Hardwick et al. 2017). Using internal standards with discriminable sequence variants enables effective normalization of inter-sample variations both in targeted RNA-seq and whole RNA-seq data, achieving accurate and reproducible quantification of gene expression (Blomquist et al. 2013; Yu et al. 2016). Therefore, constructing species-specific external RNA controls as internal standards by introducing discriminable sequence variants into endogenous transcripts can further enhance the commutability of internal standards.

Here, we designed the human-specific spike-in standards for RNA-seq experiments, systematically analyzed its technical performance, and evaluated the biases across different libraries, platforms, and laboratories. Unlike ERCC spike-in RNAs synthesized mainly from artificial sequences or non-human genes, we synthesized spike-in RNAs derived from human transcripts with artificially introduced random mutations according to the transcript length and GC content. To assess the technical performance and measure differential expression, four mixtures of these spike-in RNAs covering a 2^20^ dynamic range were obtained as spike-in standards and the RNA sample from human 293T cells was used as human reference RNA. The mixtures of these spike-in RNAs and the human reference RNA spiked with spike-in RNA mixtures were used for sequencing on Illumina and MGI platforms with different library construction methods and laboratories. We aimed to provide the human-specific spike-in standards for RNA-seq experiments and facilitate comparative analysis across different libraries, platforms, and laboratories.

## Results

### Synthesis of human-specific spike-in RNAs

Spike-in RNAs can be used as controls to evaluate the technical performance of RNA-seq experiments. Human-specific spike-in RNAs can mimic human endogenous transcripts with similar properties. Therefore, we constructed human-specific spike-in RNAs by introducing mutation sites based on human transcript sequences. Due to the longer length and higher GC content of human transcripts, we synthesized 65 human-specific spike-in RNAs with an interquartile range (IQR) of 1,590–4,032 nt in length and 44%–59% in GC content by *in vitro* transcription (**Figure 1A,B**). We evaluated the effect of mutation sites on the secondary structure of endogenous transcripts and found that the changes in minimum free energy (MFE) before and after mutation were minimal, indicating a high degree of structural similarity (**Figure 1C,D**). We found that the maximum change in minimum free energy before and after mutations was 7.2 kJ/mol, with the secondary structures revealing significant similarity before and after mutation (Similarity score: 7.84, P-value = 2.18e-06) (**Supplementary Figure S1**).

**Figure 1.**
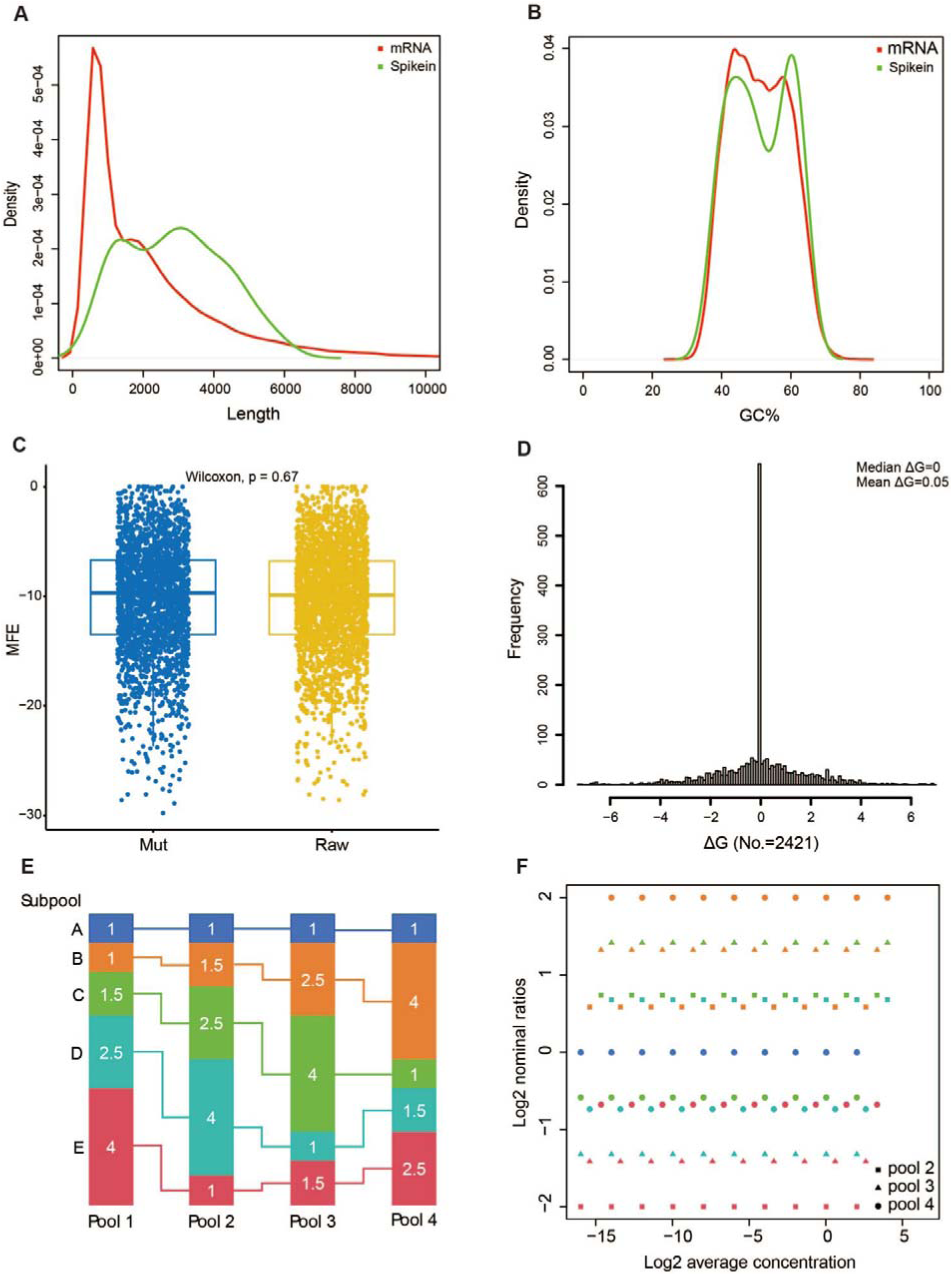
Design of spike-in RNAs and their ratio pools. **A** and **B.** Length and GC content distribution between human mRNAs and spike-in RNAs. **C.** Comparison of minimum free energy (MFE) for all mutation site regions before and after substitution. **D.** MFE change (ΔG) for all mutation site regions before and after substitution. **E.** 65 spike-in RNAs were mixed into four ratio pools composed of five subpools with 13 spike-in RNAs per subpool. **F.** Nominal ratios and abundance ranges of these spike-in RNAs (three points overlap in each subpool) compared other pools to pool 1.

### Design and evaluation of RNA-seq experiments

First, based on the concentration of spike-in RNAs, we calculated their copy number per microliter (copies/μl) and achieved the desired concentration through dilution. As a result, four spike-in mixtures/pools with 2^20^ relative abundance range were obtained to assess RNA-seq experiments (**Figure 1E,F**, and **Supplementary Table S1**). We measured the concentration of these spike-in RNAs through two rounds of digital polymerase chain reaction (ddPCR) experiments. The correlation coefficient between the two ddPCR experiments was 0.9439∼0.9928. However, significant differences were observed for the spike-in RNA pSIRC-44 and pSIRC-80 (pSIRC-44: ddPCR1/ddPCR2 = 8.93/1; pSIRC-80: ddPCR1/ddPCR2 = 1.80/1). After excluding pSIRC-44 and pSIRC-80, the correlation coefficient between the two ddPCR experiments was 0.9954∼0.9971 (**Figure 2A,B**). The concentrations in the ddPCR1 experiment were used for downstream analysis. Then, RNA-seq experiments were designed from the following aspects in this study, including four spike-in RNA mixtures and 293T RNA spiked with these mixtures (pool 1, 2, 3 and 4), three library construction protocols (rRNA-depleted, polyA(+)-selected, and 100% spike-in RNAs, assigned as RR, OLT, and SPI), two laboratories (Novogene and BerryGenomics), two sequencing platforms (Illumina and MGI, assigned as I and D), and three replicate libraries (replicate 1, 2 and 3). The laboratory and sequencing platform were combined, designating Illumina and MGI platforms at Novogene as I and D, and Illumina platform at BerryGenomics as SI. For example, the RNA-seq experiments conducted by the BerryGenomics laboratory using the Illumina platform involved sequencing three replicate libraries constructed by rRNA-depleted method from the human reference RNA spiked with spike-in RNA pool 1, designated as SI-RRpool1-1, SI-RRpool1-2, and SI-RRpool1-3 (**Figure 2C**).

**Figure 2.**
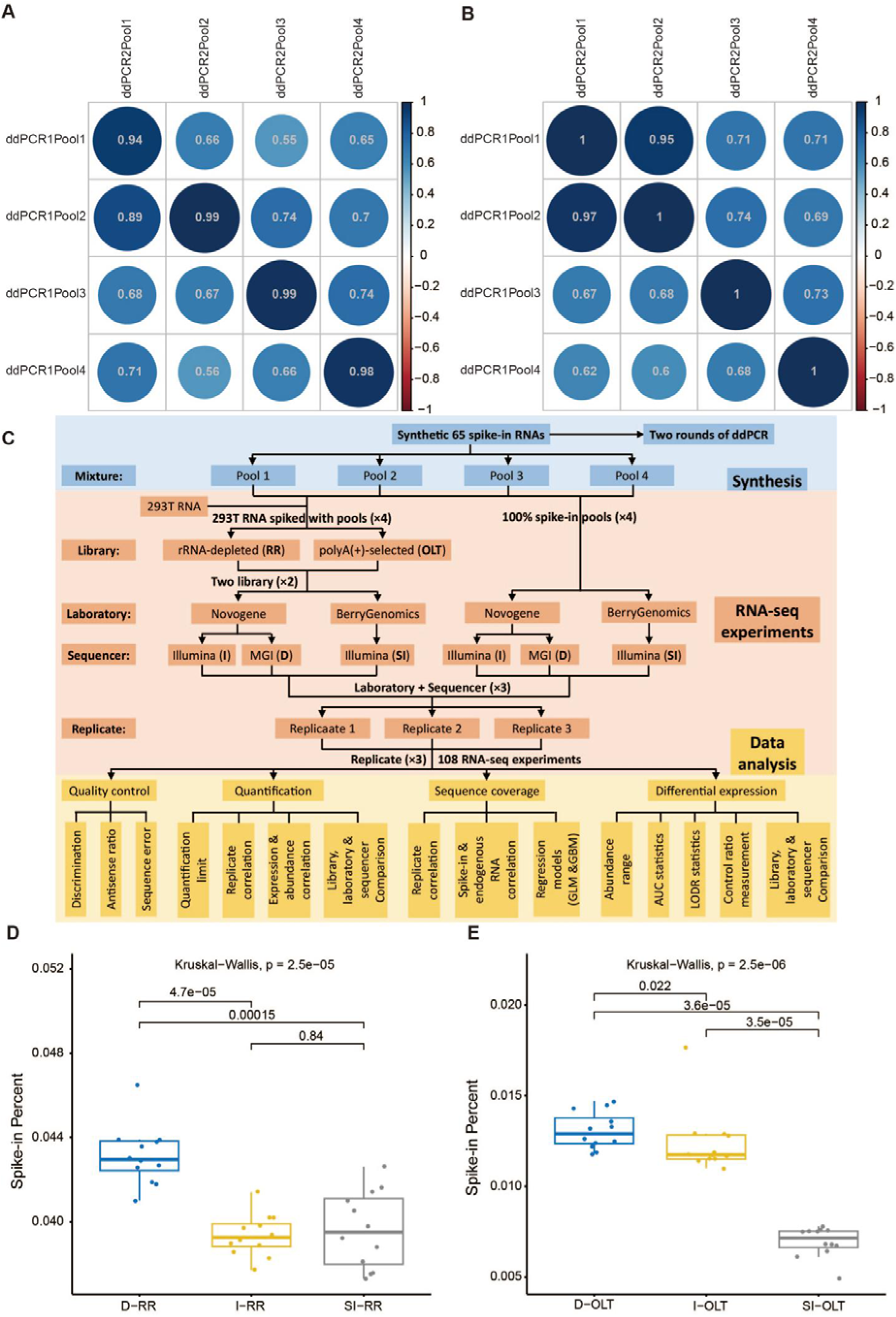
Characteristics of the spike-in RNAs. **A** and **B.** Concentration correlation of spike-in RNA pools between Droplet Digital PCR 1 (ddPCR1) and ddPCR2. All 65 spike-in RNAs were used in **A**, but 63 spike-in RNAs were used in **B** after excluding pSIRC-44 and pSIRC-80 with significant discrepancies between ddPCR1 and ddPCR2. **C.** Overview of synthesis, RNA-seq experiments and data analysis workflow. **D** and **E.** Spike-in RNA read percents in rRNA-depleted (**D**) and poly(A+)-selected (**E**) libraries, respectively.

Based on the experimental design, we constructed 108 sequencing libraries and obtained their data. Our analysis revealed that 100% spike-in RNA libraries had an average alignment rate of less than 0.008% to the human genome, indicating minimal confounding alignment between spike-in RNAs and human genome, thereby ensuring the reliability of subsequent data analysis (**Supplementary Table S2**). Using the human reference RNA spiked with 0.2% (wt/wt) spike-in RNA mixtures for RNA sequencing, we found that spike-in RNA reads accounted for only ∼1.08% of the total reads in OLT libraries, whereas RR libraries demonstrated a higher proportion (∼4.07%) (**Figure 2D,E**).

The use of spike-in RNAs enables quality control in RNA-seq experiments, including quantification of strand-specificity fidelity and systematic sequencing error rates. Data analysis showed that the proportion of antisense transcripts was approximately 1.91% (±1.57%), which exhibited potential laboratory-specific variations (**Figure 3A-C**). For instance, the BerryGenomics laboratory revealed minimal variation in antisense transcript proportions (∼1.27%) across different library construction methods, whereas substantial discrepancies were observed among distinct library preparation methods in the Novogene laboratory (0.26%, 4.51% and 1.94% in SPI, RR and OLT libraries, respectively) (**Supplementary Table S3**). Moreover, correlation analysis of spike-in RNAs revealed positive correlations between sense/antisense expression and concentration (R = 0.9891 and P = 0 for sense and R = 0.8698 and P = 0 for antisense in SI-SPIpool4-3), while a significant positive correlation was also observed between sense and antisense expression (R = 0.8514 and P = 0, SI-SPIpool4-3) (**Supplementary Figure S2**).

**Figure 3.**
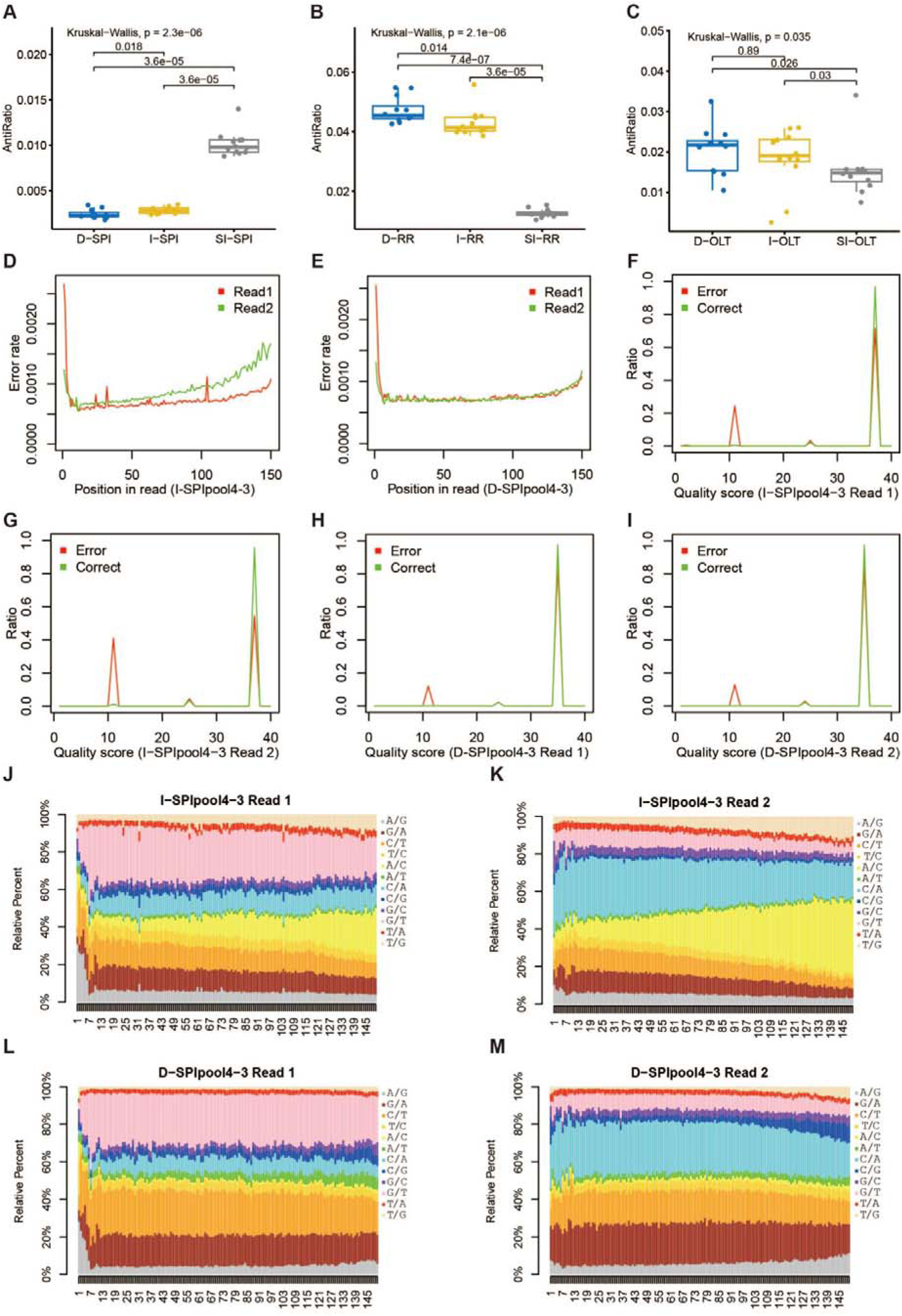
Quality control of RNA-seq experiments using the spike-in RNAs. **A**, **B** and **C.** Proportion of antisense reads in 100% spike-in RNAs (**A**), rRNA-depleted (**B**), and poly(A+)-selected (**C**) libraries. **D** and **E.** Sequencing error rate along spike-in RNA reads in I-SPIpool4-3 (**D**) and D-SPIpool4-3 (**E**). **F** and **G.** Base quality scores in reads 1 (**F**) and 2 (**G**) of spike-in RNAs in I-SPIpool4-3. **H** and **I**. Base quality scores in reads 1 (**H**) and 2 (**I**) of spike-in RNAs in D-SPIpool4-3. **J** and **K**. True-to-erroneous base change profiles in reads 1 (**J**) and 2 (**K**) of spike-in RNAs in I-SPIpool4-3. **L** and **M.** True-to-erroneous base change profiles in reads 1 (**L**) and 2 (**M**) of spike-in RNAs in D-SPIpool4-3.

As previous study, we observed that the first six nucleotides of read one, corresponding to the random hexamer priming sites during the reverse transcriptase reaction in library preparation, exhibited the highest sequence error rates with decreased tendency (**Figure 3D,E**) (Jiang et al. 2011). The error rates at the most of remaining positions were comparable until they began to increase gradually at the reads end. Moreover, excluding the first six nucleotides, read two showed consistently higher sequencing error rates than read one on Illumina platform, whereas similar error rates were observed between read one and two on MGI platform. Similarly, read two exhibited a higher proportion of low-quality scores in sequencing error sites compared to read one on Illumina platform, whereas comparable proportions of low-quality scores were noted between read one and two on MGI platform (**Figure 3F-I**). Furthermore, we identified sequencing error biases for true-to-erroneous base changes. Six nucleotides corresponding to the random hexamer priming sites preferred the A->G and C->T changes. With an increasing number of sequencing cycles, Illumina platform showed progressive accumulation of A->C and T->G changes, whereas MGI platform exhibited a gradual increase in A->G, C->G, and T->G changes. Additionally, distinct dominant true-to-erroneous base change profiles were observed between two sequencing platforms (**Figure 3J-M**).

### Quantification standard curves

Transcript quantification is critical for transcriptome studies, and spike-in RNAs serve to directly establish a quantitative correlation between transcript concentration and expression. For the 100% spike-in RNA libraries, all spike-in RNAs were detected in RNA-seq experiments, except for pSIRC-45, which failed in pool 1 and pool 4 of the SI-SPI RNA-seq experiments. Based on the concentration, pSIRC-61, pSIRC-65, and pSIRC-45 were below the quantification limit and were three of the four lowest expression spike-in RNAs in pool 1, and pSIRC-61 and pSIRC-45 were below the quantification limit and were two of the three lowest expression spike-in RNAs in pool 4.

After filtering out spike-in RNAs below the quantification limit, we conducted correlation analysis of RNA expression among replicate RNA-seq libraries as well as correlation analysis between RNA expression and concentration. First, a high degree of correlation was observed between replicates of SI-SPI, SI-RR, and SI-OLT RNA-seq libraries in spike-in RNAs (on average, R = 0.9990 on normal scale) and human reference RNAs (on average, R = 0.9974 on normal scale) (**Figure 4A-D** and **Supplementary Table S4**). Next, we built the quantification standard curve models based on spike-in RNA concentration and expression. We selected TPM value to measure RNA expression because it can effectively eliminate statistical biases for RNA length and sequencing depth of a sample (Wagner et al. 2012). The components related with RNA expression, such as RNA concentration, length, GC content, coefficient of variation (CV) for GC content, MFE, and CV for MFE, were used as the independent variables to construct the relationship with the RNA expression (TPM value) (**see Methods**). Although RNA length, CV for GC content, and CV for MFE contributed significantly to the models, only RNA concentration was used in the final model because it had the lowest Bayesian Information Criterion (BIC) values.

**Figure 4.**
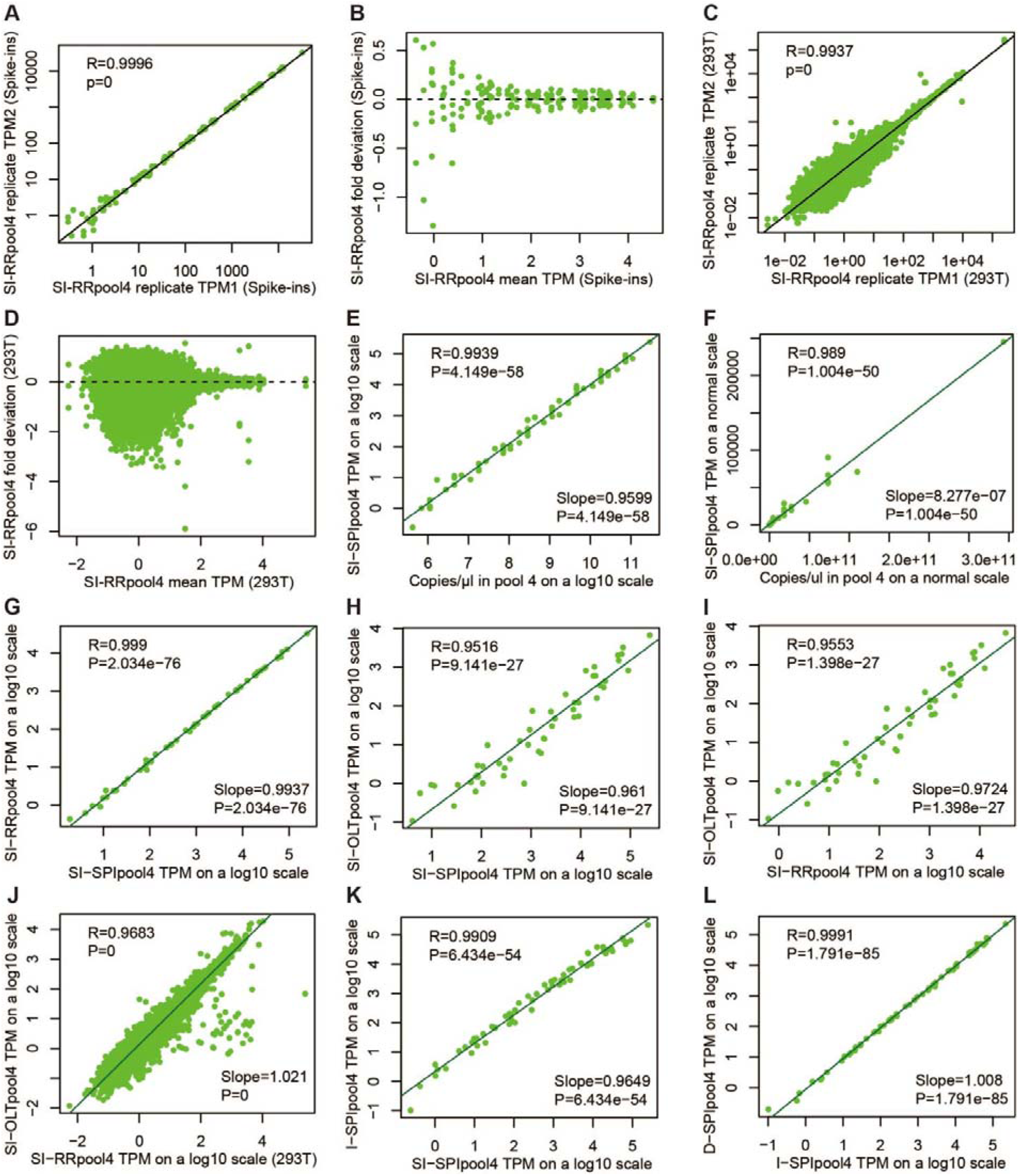
Correlation and quantification standard curve analysis. **A.** Correlation of spike-in RNA expression between replicate libraries in SI-RRpool4. **B.** Scatter plot of fold deviation between TPM in each replicate and average TPM of three replicates for each spike-in RNA in SI-RRpool4. Fold deviation is on a log2 scale and mean TPM is on a log10 scale. **C.** Correlation of 293T RNA expression between replicate libraries in SI-RRpool4. **D.** Scatter plot of fold deviation between TPM in each replicate and average TPM of three replicates for each 293T RNA in SI-RRpool4. Fold deviation is on a log2 scale and mean TPM is on a log10 scale. **E** and **F.** Correlation between spike-in RNA expression (TPM) and concentration (copies/ul) on a log10 scale (**E**) and normal scale (**F**) in SI-SPIpool4. **G**, **H** and **I.** Correlation of spike-in RNA expression (TPM) between SI-SPIpool4 and SI-RRpool4 (**G**), SI-SPIpool4 and SI-OLTpool4 (**H**), and SI-RRpool4 and SI-OLTpool4 (**I**) on a log10 scale. **J.** Correlation of 293T RNA expression (TPM) between SI-RRpool4 and SI-OLTpool4 on a log10 scale. **K** and **L.** Correlation of spike-in RNA expression (TPM) between different laboratories (**K**) and sequencing platforms (**L**) on a log10 scale.

Based on the quantification standard curve model and correlation analysis, a strong relationship was observed between RNA expression and concentration. For SI-SPI RNA-seq experiments (100% spike-in RNA libraries), the average Pearson correlation coefficient was 0.9940 (R ≥ 0.9922) on a log scale and 0.9881 (R ≥ 0.9829) on a normal scale (**Figure 4E,F** and **Supplementary Table S5**). We also achieved high correlation for spiked human reference RNA in SI-RR (on average, R = 0.9912 on a log scale and R = 0.9898 on a normal scale) and SI-OLT (on average, R = 0.9740 on a log scale and R = 0.9703 on a normal scale) RNA-seq experiments. The similar results were obtained in all other RNA-seq experiments (**Supplementary Figure S3**). Compared to R value on a normal scale, R value on a log scale slightly increased (median:1.30%, IQR: 0.38%–3.28%). We also inspected the slopes of quantification standard curve models on a log scale and found that the median slope was 0.9626 with IQR of 0.9589–0.9666, 0.9546 with IQR of 0.9405–0.96764, and 1.0556 with IQR of 1.0368–1.0770 in SI-SPI, SI-RR, and SI-OLT RNA-seq experiments, respectively (**Supplementary Table S5**). The results revealed that the model slopes for RNA-seq experiments using SPI and RR library preparation protocols were less than 1, whereas those for RNA-seq experiments using OLT library preparation protocol were greater than 1. Similar differences in slopes were observed in all other RNA-seq experiments, reflecting the possible impact of library construction (**Supplementary Figure S4**). Compared with the correlation between RNA expression and concentration, RNA expression of replicate RNA-seq libraries had a better correlation, suggesting systematic biases in RNA-seq experiments.

Moreover, the expression of spike-in RNAs was significantly correlated among SPI, RR and OLT libraries (on average on a log scale, R = 0.9892 between SPI and RR, R = 0.9583 between SPI and OLT and R = 0.9565 between RR and OLT) (**Figure 4G-I** and **Supplementary Table S6**). We also found that human transcript expression was significantly correlated between RR and OLT libraries on a log scale (on average, R = 0.9665) (**Figure 4J** and **Supplementary Table S7**). The above results indicated that spike-in RNA expression is minimally influenced by endogenous RNAs (<3% between SPI and RR (non-rRNA) libraries), but marginally effected by library preparation methods (3%–5% among non-rRNA and OLT libraries). Unlike library construction, we found that spike-in RNA expression was highly correlated between different laboratories (on average on a log scale, R = 0.9935 between SI and I) (**Figure 4K**), sequencing platforms (on average on a log scale, R = 0.9986 between I and D) (**Figure 4L**), or combinations of laboratories and sequencing platforms (on average on a log scale, R = 0.9928 between SI and D) (**Supplementary Figure S5**).

### Sequence coverage profiles

The uniformity of sequence coverage significantly affects RNA expression. These spike-in RNAs can be used to study sequence coverage and its influencing factors in RNA-seq experiments. We first obtained the sequence coverage of spike-in RNAs and their corresponding endogenous transcripts calculated according to positions and 100 equal bins across the full-length RNA. After filtering out RNAs with an average coverage of less than 1× in the three replicate libraries, the correlation of sequence coverage between different replicates was 0.9928-0.9956, 0.9777-0.9894, and 0.6686-0.8907 for all spike-in RNAs in SI-SPI, SI-RR, and SI-OLT, and 0.9034-0.9291, and 0.1362-0.6891 for all corresponding endogenous transcripts in SI-RR and SI-OLT (**Supplementary Table S8a**). The above results indicated a high degree of reproducibility of sequence coverage between replicates both on spike-in RNAs and endogenous transcripts, which have been reported in the previous studies (Li et al. 2010; Jiang et al. 2011). Moreover, the sequence coverage correlation between spike-in RNAs and corresponding endogenous transcripts using average coverage in three replicates was 0.8347 and 0.7738 in the SI-RR and SI-OLT libraries, respectively (**Supplementary Figure S6** and **Supplementary Table S8b**). However, sequence coverage of transcripts was not uniformly distributed, such as rapidly rising 5’ end, slowly descending 3’ end, and jagged region between the two (**Figure 5A-F** and **Supplementary Figure S7**). The above result indicated that library construction and local sequences contribute to non-uniform sequence coverage.

**Figure 5.**
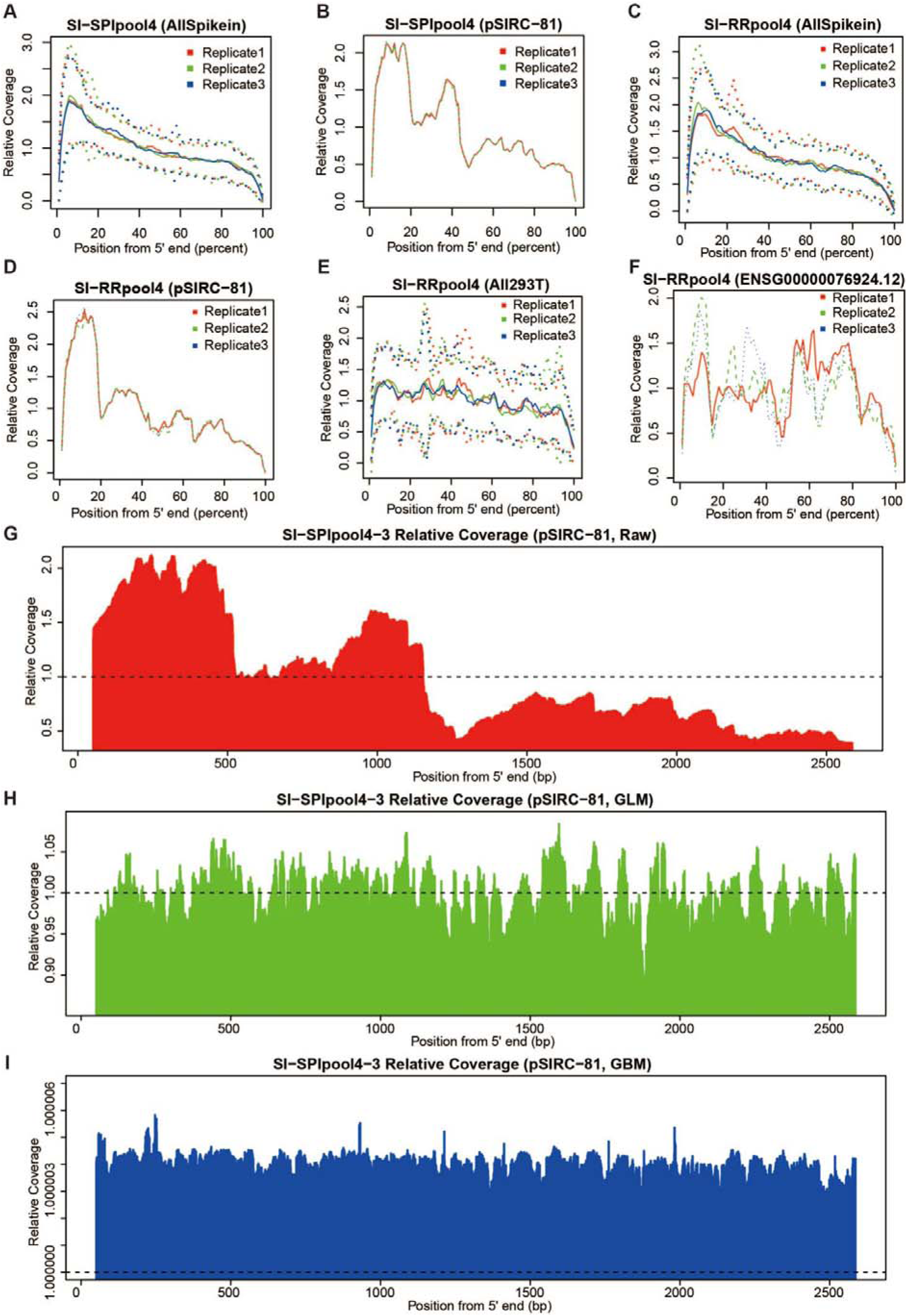
Relative coverage heterogeneity and fitting coverage. **A** and **B.** Relative coverage along all spike-in RNAs (**A**) and spike-in RNA pSIRC-81 (**B**) in SI-SPIpool4. **C** and **D.** Relative coverage along all spike-in RNAs (**C**) and spike-in RNA pSIRC-81 (**D**) in SI-RRpool4. **E** and **F.** Relative coverage along all endogenous counterparts of spike-in RNAs (**E**) and endogenous transcript of ENSG00000076924.12 (XAB2) (**F**) in SI-RRpool4. **G**, **H** and **I.** Sequence coverage along spike-in RNA pSIRC-81 in raw count (**G**), fitted count using the Poisson general linear regression model (GLM) (**H**), and fitted count using the gradient boosting regression model (GBM) (**I**).

Without considering experimental related factors, heterogeneity of sequence coverage may be related to sequence related factors, such as RNA secondary structure and GC content. Based on surrounding sequences of each position along transcripts, Poisson general linear regression model (GLM) and multiple additive regression trees model (MART) were used to smooth heterogeneity of sequence coverage in previous studies, which explained about 50% and 70% of variation in sequence coverage (Li et al. 2010; Jiang et al. 2011). However, these models only use local sequences and do not consider the RNA secondary structure and GC content. Here, we used GLM and gradient boosting regression model (GBM) to reduce sequence coverage heterogeneity, in which the read coverage of each position along transcripts was modeled as a function of expression abundance (average coverage), GC content bias (GC content and CV for GC content), and RNA secondary structure bias (MFE and CV for MFE). For our SI-SPI RNA-seq experiments (pool 3 and 4), GLM and GBM explained 86.62% to 91.78% and 87.00% to 91.49% of variation in sequence coverage based on cross-validation R^2^ and test-dataset R^2^, respectively (**Figure 5G-I**, **Supplementary Figure S8**, **Supplementary Figure S9**, and **Supplementary Table S9**). Moreover, the variance-to-mean ratio (VMR) and CV were used to measure the dispersion of sequence coverage distribution of spike-in RNAs (**Supplementary Table S10a**). All spike-in RNAs had significantly higher VMR and CV scores in raw sequence coverage than those in corrected sequence coverage by GLM and GBM models. In addition, comparative evaluation of the regression models demonstrated comparable predictive efficacy between GLM and GBM based on standardized performance metrics (Mean Relative Error: GLM=0.5832±0.0087 *vs.* GBM=0.5965±0.0170; R²=0.8925 *vs.* 0.8918) (**Supplementary Table S9** and **Supplementary Table S10b**).

### Differential gene expression

The number of sequencing fragments (25.63 million fragments or 7.69 Gb in average) was more or less the same in different libraries (25.87, 26.27 and 24.76 million fragments for RR, OLT and SPI libraries) (**Supplementary Table 2**). Based on the limit of detection (LOD), the range of RNA abundance on average in SPI and RR libraries was 2^20^, while 2^18^ in OLT library (**Figure 6A**, **Supplementary Figure S10A**, **Supplementary Figure S11A**, and **Supplementary Table S11**). This difference was mainly because of decreased spike-in RNA input (100% (wt/wt), 0.2% (wt/wt) and 0.2% (wt/wt) spike-in RNA input for SPI, RR and OLT libraries) and spike-in RNA fragment ratio (96.92%, 4.07% and 1.08% spike-in RNA fragments for SPI, RR and OLT libraries) in these RNA-seq libraries. According to the limit of quantification (LOQ), the RNA abundance range on average was 2^20^, 2^17^, and 2^15^ in SPI, RR and OLT libraries, respectively (**Figure 6B** and **Supplementary Table S11**).

**Figure 6.**
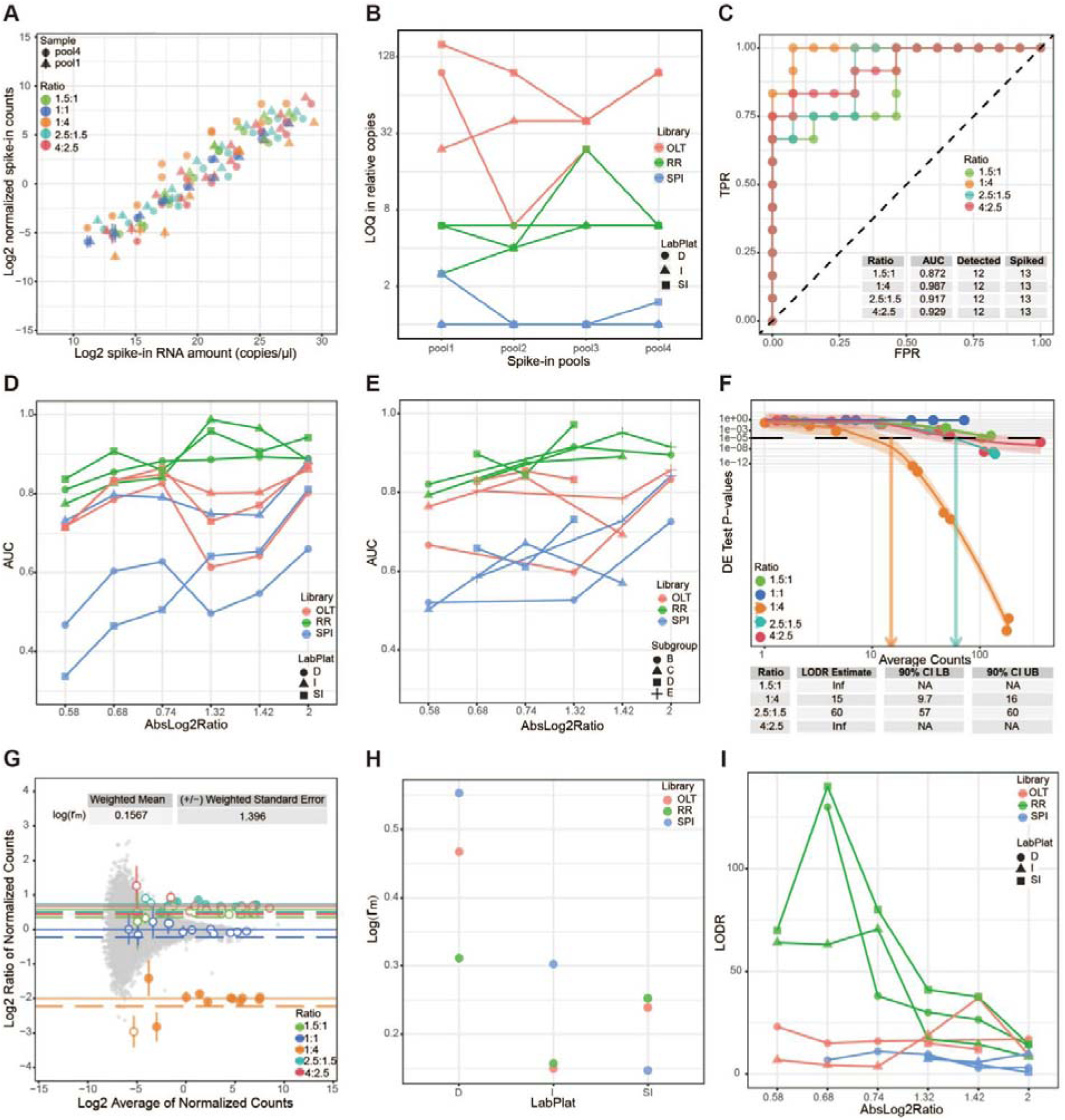
Technical performance in differential expression with spike-in RNA mixtures. **A.** Signal-abundance plot showed the dynamic range of SI-RRpool1 and SI-RRpool4 RNA-seq experiments with technical replicates (n=3). Each point was colored by ratio between two subpools, error bars represented the standard deviation of replicates, and shape represented samples. **B.** Limit of quantification (LOQ) in relative copies of spike-in RNAs in all samples. **C.** Receiver operator characteristic (ROC) curves and area under curve (AUC) statistics between SI-RRpool1 and SI-RRpool4 RNA-seq experiments. **D** and **E.** AUC statistics of these spike-in RNAs compared other pools to pool 1 at six different ratios in all samples. **D** was grouped by library in color and laboratory-platform (LabPlat) in shape, and **E** was grouped by library in color and subpool in shape. **F.** Limit of detection of ratio (LODR) estimate using erccdashboard R package between SI-RRpool1 and SI-RRpool4 RNA-seq experiments. The black dashed line annotates the P-value threshold derived from the FDR chosen (0.05). Colored arrows indicate the LODR estimate (average counts) for each ratio. **G.** MA plot showed control ratio variability and bias between SI-RRpool1 and SI-RRpool4 RNA-seq experiments. Spike-in RNA points (colored by ratio) represented the mean ratio measurements. Error bars represent the standard deviation of replicate ratios. Filled circles indicate ratios above the LODR estimate. Endogenous transcript ratio measurements are shown as grey points. The nominal ratios are annotated with solid lines for each subpool and the adjusted ratios are annotated with dashed lines. **H.** Log(r_m_) was grouped by library in color. **I.** LODR estimates were grouped by library in color and laboratory-platform (LabPlat) in shape.

The defined abundance ratios among different spike-in RNA mixtures (**Figure 1E,F**) can be used to assess the technical performance of differential gene expression. Based on ratio mixture analysis with erccdashboard package (Munro et al. 2014), all area under the curve (AUC) statistics were over 0.81 (median:0.94) with absolute log2foldchange > 1 in the RR RNA-seq experiments (**Supplementary Table S12**). The AUC statistics improved with the increase of abundance ratio and expression abundance of spike-in RNAs in general (**Figure 6C,D**, **Supplementary Figure S10B**, **Supplementary Figure S11B**). Among the four subgroups with different abundance ratios, subgroups B and C had a lower AUC in most cases compared with other subgroups, indicating differences in differential expression among these subgroups (**Figure 6E**). Moreover, AUC value is higher in RR RNA-seq experiments than that in OLT RNA-seq experiments, which probably be related with the number of low expression spike-in RNAs (expressed spike-in RNAs below LOQ: 4.92 *vs.* 8.17 in average for RR *vs.* OLT, P = 1.1E−02, *t*-test). The SPI libraries have the lowest AUC statistics due to only spike-in RNAs in these libraries (**Supplementary Table S12**).

As the above results in AUC statistics, the estimates of limit of detection of ratio (LODR) showed that the performance improved with expression abundance and abundance ratio in general (**Figure 6F**, **Supplementary Figure S10C**, and **Supplementary Figure S11C**). The bias and variability in control ratio measurements were evaluated by log value of control ratio (log(r_m_)) and its weighted standard error, and illustrated in MA plot (**Figure 6G**, **Supplementary Figure S10D**, and **Supplementary Figure S11D**). We found that the fluctuation of control ratio is gradually decreased with the increase of expression abundance. Moreover, log(r_m_) value seems to be irregular in different RNA-seq experiments (**Figure 6H**).

Three different aspects of RNA-seq experiments including library construction, sequencing platform and different laboratory were compared in this study. The AUC statistics improved with increase of abundance ratios of spike-in RNAs and they were higher in RR than those in OLT and SPI RNA-seq experiments (**Figure 6D**). Compared with MGI platform at Novogene, Illumina platforms at Novogene and BerryGenomics had better AUC performance when the absolute log2foldchange of abundance ratios was over 1. Similarly, LODR estimates also showed better performance in RR (detected in all abundance ratios) than those in OLT (detected in 9 of 18 abundance ratios) and SPI (detected in 8 of 18 abundance ratios) RNA-seq experiments when the absolute log2foldchange of abundance ratios was greater than 1 (**Figure 6I** and **Supplementary Table S12**). Moreover, Illumina platform at Novogene had better (lower) LODR estimates than those for MGI platform at Novogene and Illumina platform at BerryGenomics in RR RNA-seq experiments. Unlike AUC and LODR, log(r_m_) estimates were variable in different RNA-seq experiments (**Figure 6H**). However, Illumina platforms at Novogene and BerryGenomics had better log(r_m_) estimates than MGI platform at Novogene. The slightly worse performances of AUC, LODR and log(r_m_) probably be related to the lower LOQ on MGI platform than those on Illumina platform (LOQ: 13.33 *vs.* 18.54 in average for D *vs.* I; 13.33 *vs.* 36.50 in average for D *vs.* SI).

## Discussion

Synthetic spike-in RNAs are served as the reference standards for quantitative reverse transcriptase PCR (qRT-PCR), microarray, and RNA-seq assays (Hardwick et al. 2017). Although external spike-in RNAs have been developed to mimic eukaryotic transcripts as closely as possible, most of which were either artificially synthesized or derived from non-mammalian species (External 2005). Constructing species-specific spike-in RNAs by introducing sequence variants into endogenous transcripts can enhance commutability. In this study, a library of 65 human-specific spike-in RNAs was developed by introducing mutations into endogenous transcripts and assessed as the external RNA reference standards for bulk RNA-seq experiments. More importantly, each of our spike-in RNAs was paired with a corresponding endogenous transcript, enabling detailed comparative analysis of these transcriptions.

Despite being the most widely utilized spike-in controls, persistent discrepancies remain between ERCC controls and human transcripts. In comparison to human transcripts, which exhibit an IQR of 709–2,881 nucleotides (nt) in length and 44%–58% in GC content, ERCC controls demonstrate a shorter length (median: 994 nt, IQR: 541–1,060 nt) and a reduce GC content (median: 49%, IQR: 38%–51%) (Lee et al. 2016). To align with the long length and high GC content of human transcripts, our spike-in RNAs present optimized characteristics with an IQR of 1,590–4,032 nt in length and 44%–59% in GC content. Moreover, our synthesized human-specific spike-in RNAs demonstrate high similarity with endogenous transcripts based on the comparative analysis of MFEs and RNA secondary structures.

Using the synthesized spike-in RNAs, we constructed four mixed pools using a modified Latin square design. A total of 108 RNA-seq experiments were carried out for four 100% spike-in RNA pools and 293T cell RNA spiked with 0.2% (wt/wt) spike-in RNA pools, and their performance was systematically assessed across three library preparation protocols (SPI, RR and OLT), two sequencing platforms (Illumina and MGI), and two independent laboratories (Novogene and BerryGenomics) with three experimental replicates. Although our human-specific spike-in RNAs were highly similar to their endogenous counterparts, the average alignment rate to the human genome was <0.008% on 100% spike-in RNA libraries, which is slightly lower than the genomic alignment proportion (<0.01%) in the human genome observed in the 100% ERCC library (Jiang et al. 2011). The above result showed that synthesized spike-in RNAs are discriminable from human endogenous transcripts, indicating their use as spike-in standards in human for RNA-seq experiments. Based on alignment statistics, the proportion of spike-in RNA reads was different for OLT and RR libraries (1.08% for OLT libraries *vs.* 4.07% for RR libraries). The lower proportion of spike-in RNA reads in the OLT libraries might be attributed to their shorter polyA(+) tails (20 nt), which exhibit reduced binding efficiency during polyA(+) selection. In contrast, rRNA depletion preserves more intact polyA(+) RNA. Previous studies indicated that the mapping ratio of ERCC spike-in RNAs varies widely in library constructions (Qing et al. 2013; Consortium 2014; Risso et al. 2014). Considering the detection and quantification limit of spike-in RNAs, we recommend that 0.2%–0.4% (wt/wt) and 0.8%–1.6% (wt/wt) spike-in RNA mixture should be added in human target RNA for RR and OLT libraries.

A previous large-scale study failed to capture the overall sequencing error landscape because it relied on overlapping regions of paired-end reads to identify sequencing errors (Stoler and Nekrutenko 2021). Through systematic analysis of true-to-erroneous base change profiles, we obtained the following results: 1) unlike the Illumina platform, the sequencing error rates and base quality scores on the MGI platform were similar between reads 1 and 2; 2) the true-to-erroneous base change profiles differed between the two platforms; the Illumina platform showed gradual accumulation of A→C and T→G changes, whereas the MGI platform exhibited progressive increases in A→G, C→G, and T→G changes with increasing sequencing cycles; and 3) the sequencing error signatures were distinct between the two platforms.

In our study, slope differentials were observed between the library construction methods based on quantitative standard curve analysis (0.941 *vs.* 1.055 in average for SPI and OLT libraries, P = 7.396e-07, Wilcoxon test; 0.942 *vs.* 1.055 in average for RR and OLT libraries, P = 7.396e-07, Wilcoxon test), demonstrating the necessity of library-specific normalization during cross-library transcriptome data integration. The previous study showed that there is a strong library preparation effect although a good linear relationship between reads counts and concentrations has been observed (Risso et al. 2014). Moreover, the discrepancy in spike-in RNA quantification was also noticed between RR and OLT libraries (R = 0.9565), mechanistically linked to their intrinsic transcript selection methods: rRNA depletion and polyA(+) selection. Furthermore, the technical discrepancy between RR and OLT libraries was further confirmed by differential coverage distribution (5’-to-3’ coverage gradient in RR libraries *vs.* relatively uniform coverage in OLT libraries).

The sequence coverage trends of transcripts are reproducible, yet exhibit marked heterogeneity. The sequence coverage correlation between spike-in RNAs and corresponding endogenous transcripts was compared for the first time and the correlation coefficient was high (R = 0.8347 and R = 0.7738 in SI-RR and SI-OLT libraries, respectively). Previous studies constructed regression models based on the primary sequence features of transcripts, accounting for 50% to 70% of the variation in sequence coverage (Li et al. 2010; Jiang et al. 2011). Based on RNA secondary structure and GC content of local sequence, we built GLM and GBM models, which accounted for 86.62%–91.78% and 87.00%–91.49% of the variation in sequence coverage, respectively. Compared to the previous studies, the R^2^ values increased more than 20% in our models.

We evaluated the technical performance of differential gene expression in RNA-seq experiments and observed bias contributed by library preparation procedures as before (Munro et al. 2014). According to the AUC statistics and LODR estimates, differential expression identification is more accurate for genes with an fold change of ≥2 between two samples and library construction methods have more impact than laboratories and sequencing platforms. The median AUC values were 0.94, 0.80, and 0.72 with an absolute log2foldchange > 1 in RR, OLT, and SPI RNA-seq experiments, respectively. The RR RNA-seq experiments achieved better AUC statistics compared to OLT (increased with an median of 0.140 and IQR of 0.119-0.189) and SPI (increased with an median of 0.230 and IQR of 0.193-0.265) RNA-seq experiments. For laboratories, Illumina platform at Novogene acquired higher AUC statistics than Illumina platform at BerryGenomics (increased with an median of 0.060 and IQR of 0.017-0.082). For sequencing platforms, the AUC values increased with an median of 0.100 and IQR of 0.045-0.266 comparing Illumina with MGI platform at Novogene. Similarly, RR RNA-seq experiments obtained all LODR estimates, while equal to or more than half of OLT and SPI RNA-seq experiments did not obtain LODR estimates when the absolute log2foldchange of abundance ratios was greater than 1. The similar results were obtained in different laboratories and sequencing platforms for RNA-seq experiments between LODR estimates and AUC statistics. Moreover, log(r_m_) values exhibited irregular fluctuations, which may be influenced by other uncertain factors except for library construction, sequencing platform and laboratory.

Our study has several limitations. First, we only synthesized 65 human-specific spike-in RNAs, which represents a limited number compared to the ERCC spike-in controls. Second, only 293T cells were used and prevented the evaluation across different cell lines or tissues in RNA-seq experiments. Third, human 293T cell RNA spiked with only 0.2% (wt/wt) spike-in RNAs was used for RNA-seq experiments. We propose that adding 0.2%–0.4% (wt/wt) and 0.8%–1.6% (wt/wt) spike-in RNAs to RR and OLT libraries, respectively, would enable the detection of most spike-in RNAs. Finally, we incorporated a 16-nt barcode at the 5’ end of all spike-in RNAs to serve as identifier sequences for third-generation sequencing, but we did not carry out any work on long-read RNA-seq experiments. Nevertheless, we developed a set of human-specific spike-in RNAs that enable rigorous evaluation of human RNA-seq experiments. Additionally, our study provides a basis for comparative analyses between species-specific and universal spike-in controls, which was previously unfeasible.

## Methods

### Human-specific spike-in RNA production

Preliminary RNA sequences were extracted according to the length and GC content distribution of human transcripts (GCF_000001405.39). Then, random mutations were introduced in each RNA approximately every 75 bp after excluding SNP sites in dbSNP (Phan et al. 2025). Next, the plasmid DNA libraries of these RNA sequences were constructed using similar procedure for ERCC (https://www.nist.gov/system/files/documents/2016/09/26/2374_coa_2013.pdf). Compared to ERCC, we have made the following modifications: 1) the pUC57 vector was used for plasmid construction; 2) the promoter sequence of T7 RNA polymerase and the 16 base barcode sequence were added at the 5′ end for *in vitro* transcription and unique molecular identifier; 3) a 20 base polyadenylated (polyA(+)) fragment was added at the 3′ end for mimicking eukaryotic messenger RNA (mRNA). Sanger sequencing were used to confirm the sequences of these spike-in RNAs. After that, the plasmid DNA libraries were used for PCR amplification and subsequent *in vitro* transcription using the T7 Quick High Yield RNA Synthesis Kit (New England Biolabs Inc., Ipswich, MA) (**Supplementary Figure 12**).

### Pool design

Four mixtures of these spike-in RNAs (pool 1 to pool 4) were obtained as spike-in RNA standards. Each mixture was designed to have a 2^20^ dynamic range and was composed of five subpools (13 spike-in RNAs in each subpool). The five subpools in different mixtures were present with defined abundance ratios among these mixtures according to a modified Latin square design (**Figure 1E,F**). One subpool has identical spike-in RNA abundances while the other four subpools have different abundances in the four mixtures. Moreover, the abundance ratios of five subpools in each mixture are fixed and the abundance ratios of the other four subpools to the identical abundance subpool were 1:1, 1.5:1, 2.5:1, and 4:1, respectively.

### RNA sequencing (RNA-seq)

The total RNA was extracted from human 293T cells and was used as human reference RNA in this study. Each mixture alone and human reference RNA spiked with mixture at 0.2% (wt/wt) were used for RNA sequencing. PolyA(+) RNA was extracted with VAHTS mRNA Capture Beads (Vazyme, Nanjing, China) and rRNA-depleted RNA was isolated using Ribo-off rRNA Depletion Kit (Vazyme, Nanjing, China). The dUTP protocol were used to construct strand-specific library. Three technical replicates were done for each RNA-seq experiment. For comparison between sequencing platforms and laboratories, Illumina (Novaseq 6000) and MGI (DNBSEQ-T7) platforms were used for sequencing in Novogene (Beijing, China) and Illumina platform (Novaseq 6000) was used in BerryGenomics (Beijing, China).

### Sequencing analysis

150 bp paired-end reads were produced for each sequencing sample with an average total fragments of 25.63 millions (**Supplementary Table 2**). Quality control of raw sequencing data was done using FastQC (v0.11.9) and adapter and low quality sequences were filtered using fastp (v0.20.1) with the parameters “-x −5 −3 -r -c -q 20 -u 20 -n 5 −l 100” (https://www.bioinformatics.babraham.ac.uk/projects/fastqc/) (Chen et al. 2018). Human nuclear and mitochondrial rRNAs were obtained from MINRbase as reference rRNAs and bowtie2 (v2.4.2) was used to identify rRNA reads (Langmead and Salzberg 2012; Pliatsika et al. 2024). The remained reads were aligned to spike-in RNAs and the properly aligned paired reads (mapping identity >= 98%, mapping quality >= 60, and the introduced mutation sites >= 1) were assigned as spike-in RNA reads. After removing spike-in RNA reads, the remained reads were aligned to human reference genome (The GATK resource bundle for Grch38/hg38: https://console.cloud.google.com/storage/browser/genomics-public-data/resources/broad/hg38/v0/) by HISAT2 (v2.1.0) and the properly aligned paired reads (mapping identity >= 98%, mapping quality >= 60, and without the introduced mutation site) were assigned as human reference RNA reads (Kim et al. 2019). Read counts for spike-in RNA and human reference RNA were quantified using featureCounts (v2.0.1) (Liao et al. 2014). Count data from sequencing samples were used in the downstream analysis. Transcripts Per Million (TPM) was calculated as previously described (Wagner et al. 2012). RNAfold (v2.3.3) was used to get secondary structure and minimum free energy (MFE) (Lorenz et al. 2011; Varenyk et al. 2023). GC content, CV for GC content, MFE and CV for MFE were obtained using 51 bp sliding window and 1 bp step size along whole sequences of spike-in RNAs (**Supplementary Figure 13**).

### Sequencing error analysis

Mapped spike-in RNA reads were compared with spike-in RNA sequences using an in-house Perl script to identify sequencing error bases along the reads. The types of change from the reference base to the error base were counted at each position across reads. The distribution of sequencing quality scores was calculated for both correctly sequenced and erroneously sequenced bases. Sequencing error rates were obtained by dividing the sequencing error bases by the total sequencing bases.

### Quantification standard curve models

The minimum spike-in RNA concentration with reads support in all replicates was defined as the concentration quantification limit. The spike-in RNAs exceeding the concentration quantification limit were used for model fitting. Generalized Least Squares (GLS) fitted linear model was employed using R package nlme (v3.1.163) to evaluate the linear relationship between spike-in RNA abundance related components (concentration, length, GC content, CV for GC content, MFE, and CV for MFE) and expression (TPM value). Several models were built as follows:

1. *Y* = b_0_ + b_1_*M* + b_2_*L* + b_3_*G* + b_4_*GV* + b_5_*E* + b_6_*EV* + *e* (BIC score = 25.94),
2. *Y* = b_0_ + b_1_*M* + b_2_*L* + b_4_*GV* + b_6_*EV* + *e* (BIC score = 5.83),
3. *Y* = b_0_ + b_1_*M* + b_4_*GV* + b_6_*EV* + *e* (BIC score = −6.26),
4. *Y* = b_0_ + b_1_*M* + *e* (BIC score = −6.64),

Here, *Y* stands for TPM value in RNA-seq experiment, *M* represents RNA concentration (RNA copy number), *L* is RNA length, *G* and *E* indicate GC content and MFE, *GV* and *EV* denote CV for GC content and MFE respectively. Additionally, b_0,_ b_1,_ b_2,_ b_3,_ b_4,_ b_5,_ and b_6_ are coefficients while *e* is residual error. *Y* and *M* are on a log10 scale while other components are on a normal scale. BIC is Bayesian Information Criterion.

### Sequence coverage models

The spike-in RNAs without read in any of three replicates were excluded from model fitting. General linear regression model (GLM) using R package stats (v4.3.2) and gradient boosting regression model (GBM) using R package gbm (v2.2.2) were applied to obtain sequence preference models for read coverage based on average coverage, GC content, CV for GC content, MFE and CV for MFE. The third replicate of pool 3 and pool 4 was used as the training data. A five fold cross-validation strategy was implemented to calculate R^2^ using R package caret (v6.0.94).

### Differential gene expression

The R package erccdashboard (v1.36.0) was used to evaluate the spike-in RNA ratio performance metrics for differential gene expression, which include dynamic range, diagnostic performance, LODR estimates and expression ratio bias (Munro et al. 2014).

## Data access

Raw data were deposited in the Genome Sequence Archive for human (GSA-Human: HRA010537) at the National Genomics Data Center, Beijing Institute of Genomics, Chinese Academy of Sciences / China National Center for Bioinformation, which can be accessed at https://bigd.big.ac.cn/gsa-human (Chen et al. 2021). The reviewer link of our data is https://ngdc.cncb.ac.cn/gsa-human/s/P58sIooD for HRA010537.

## Acknowledgments

This study was supported by the Biological Breeding-National Science and Technology Major Project (2022ZD04017).

## CRediT author statement

**Rui Qin**: Resources, Investigation, Formal analysis, Validation, Writing - original draft. **Wei Fan**: Methodology, Investigation, Formal analysis, Visualization. **Feng Ding**: Investigation, Formal analysis, Validation, Writing - original draft. **Sen Wang**: Investigation, Formal analysis. **Bing He**: Investigation, Formal analysis. **Mingming Hou**: Investigation, Formal analysis.**Qiang Lin**: Conceptualization, Supervision, Writing - review & editing. **Peng Cui**: Conceptualization, Supervision, Project administration, Funding acquisition, Writing - review & editing. **Wanfei Liu**: Supervision, Methodology, Investigation, Formal analysis, Visualization, Writing - original draft, Writing - review & editing. Wanfei Liu has accessed and verified the underlying data. All authors read and approved the final manuscript.

